# Neural oscillations during conditional associative learning

**DOI:** 10.1101/198838

**Authors:** Alex Clarke, Brooke M. Roberts, Charan Ranganath

## Abstract

Associative learning requires mapping between complex stimuli and behavioural responses. When multiple stimuli are involved, conditional associative learning is a gradual process with learning based on trial and error. It is established that a distributed network of regions track associative learning, however the role of neural oscillations in human learning remains less clear. Here we used scalp EEG to test how neural oscillations change during learning of arbitrary visuo-motor associations. Participants learned to associative 48 different abstract shapes to one of four button responses through trial and error over repetitions of the shapes. To quantify how well the associations were learned for each trial, we used a state-space computational model of learning that provided a probability of each trial being correct given past performance for that stimulus, that we take as a measure of the strength of the association. We used linear modelling to relate single-trial neural oscillations to single-trial measures of association strength. We found frontal midline theta oscillations during the delay period tracked learning, where theta activity was strongest during the early stages of learning and declined as the associations were formed. Further, posterior alpha and low-beta oscillations in the cue period showed strong desynchronised activity early in learning, while stronger alpha activity during the delay period were seen as associations became well learned. Moreover, the magnitude of these effects during early learning, before the associations were learned, related to improvements in memory seen on the next presentation of the stimulus. The current study provides clear evidence that frontal theta and posterior alpha/beta oscillations play a key role during associative memory formation.

## Introduction

In our daily lives, we must learn arbitrary associations between initially unrelated items, such as when we meet a specific person, in a certain place to give them a specific object. In studies using animal models, neuroscientists have used conditional associative learning paradigms as a way to study how arbitrary associations are learned (for review see Suzuki, 2008). In conditional associative learning, one must learn mappings between complex stimuli and behavioural responses, based on trial and error. Conditional associative learning is dependent on the hippocampus (HPC; Murray & Wise, 1996; Stark, Bayley, & Squire, 2002), striatum (Brasted & Wise, 2004), and a distributed network of regions in frontal and parietal cortex (Asaad, Rainer, & Miller, 1998; Law et al., 2005; Petrides, 1997) that are believed to interact with the HPC during learning (Brincat & Miller, 2015; Siapas, Lubenov, & Wilson, 2005). What has received much less focus is the role of neural oscillations in associative memory formation within this distributed network. Neural oscillations are believed to play a critical role in human cognition (Buzsáki & Draguhn, 2004; Kahana, 2006; Siegel, Donner, & Engel, 2012), and are modulated by memory performance (Hanslmayr, Staudigl, & Fellner, 2012; Hsieh & Ranganath, 2014). Here, we track how oscillatory activity changes during associative learning.

Learning multiple associations is a gradual process, and requires an approach where the strength of the association can be estimated on a trial-by-trial basis. Suzuki and colleagues have used a dynamic state-space model to estimate such learning and investigated changes in neurophysiological activity during conditional associative learning (Hargreaves, Mattfeld, Stark, & Suzuki, 2012; Law et al., 2005; Wirth et al., 2003). Such dynamic models of learning provide a probability that a particular association has been learned on a given trial, based on past behaviour, thus providing a continuous measure of association strength (Smith et al., 2004). By tracking how activity for single trials related to a single trial measure of learning, Wirth et al. (2003) found that spiking activity in the primate hippocampus showed a linear relationship with association strength, where cells either showed an increased spike rate as the association was learned, or showed a high spike rate during initial learning that reduced back to baseline levels with learning. A linear relationship between association strength and human fMRI was further observed in the MTL, frontal, temporal and parietal regions (Law et al., 2005), while beta oscillations in the monkey entorhinal cortex were shown to increase in a similar linear fashion as memories became stronger (Hargreaves et al., 2012). Outside the MTL, prefrontal oscillations in beta (Brincat & Miller, 2016) and theta frequencies (Loonis, Brincat, Antzoulatos, & Miller, 2017; Paz, Bauer, & Paré, 2008) also increase as memories are established. Together, the extant evidence suggests that a distributed network of regions tracks conditional associative learning, and recordings from nonhuman primates further indicates that theta and beta oscillations might provide a key signature of memory formation.

The role of neural oscillations in human learning, however, remains less clear. Electroencephalography (EEG) can be used to monitor oscillations generated in the human neocortex, but to our knowledge, EEG has not yet been used to test whether neural oscillations modulate the gradual learning of new associations across the human cortex. Much of what we know about the role of neural oscillations during memory formation comes from research where memories are formed from a single exposure to a stimulus, where successful memory formation is defined by whether a participant can accurately retrieve the item or not. Scalp EEG in humans has shown that theta oscillations over frontal sites increase with successful memory encoding (Hsieh & Ranganath, 2014; W. Klimesch, Doppelmayr, Schimke, & Ripper, 1997; Mölle, Marshall, Fehm, & Born, 2002; Summerfield & Mangels, 2005; White et al., 2013), while other research highlights the role of alpha and beta oscillations during memory formation (Hanslmayr, Spitzer, & Bäuml, 2009; W. Klimesch et al., 1996, 1997; Mölle et al., 2002).

In the present study, we examined how neural oscillations change during learning of arbitrary visuo-motor associations. Specifically, we investigated: (1) how oscillations change as associations are learnt and get stronger (similar to previous approaches, e.g. Hargreaves et al., 2012; Law et al., 2005; Wirth et al., 2003), and (2) how oscillations signify how much learning is taking place. We recorded human scalp EEG as participants learned to associate each of 48 abstract shapes with one of 4 button responses. By fitting a state-space model to participant behaviour, we estimated trial-by-trial estimates of association strength and related these measures to oscillations, providing a powerful and sensitive approach to understanding the role of neural oscillations during memory formation.

## Methods

### Participants, Stimuli and Procedure

Eighteen right-handed subjects took part in the study (range 18-25 years). All subjects had normal, or corrected to normal, vision and gave written informed consent prior to the study. The study was approved by the Institutional Review Board at the University of California, Davis. Two subjects performed poorly on the final test (<65% correct) and were excluded from all analyses.

Subjects performed an associative learning task where they learned to associate one of 4 button presses with an abstract shape over many repetitions of the item (Figure 1A). The items were 48 abstract shapes that were colored red, green, blue or yellow. All items were centrally positioned on a black background. Each trial began with a blank screen for 2 seconds, followed by a cue item for 1.5 seconds. A 3 seconds delay period followed, after which was a 3 second response period where the response options were displayed on screen (1, 2, 3, 4). Finally, a feedback screen informed the subject if the response they made was correct or incorrect. Each shape was repeated 12 times within a block, and each block contained 12 different shapes. All shapes were shown across 4 different blocks. There was no association between the colors and the correct responses. After the 4 blocks, a final test block was conducted where all 48 shapes were shown.

### EEG recording

EEG was recorded using a BioSemi (http://www.biosemi.com) Active Two system at a sampling rate of 2048 Hz in a sound-attenuated chamber. Recordings were made from 64 active Ag/AgCl scalp electrodes embedded in an elastic cap, with electrode locations corresponding to an extended version of the international 10/20 system. Additional recordings were made from electrodes placed on the left and right mastoids, and around the eyes (lateral to each eye, and above and below the left eye). EEG was recorded with respect to a common mode sense active electrode located on the scalp near electrode site Cz. Subjects were instructed to minimize muscle tension, eye movements and blinking during the study.

**Figure 1:**
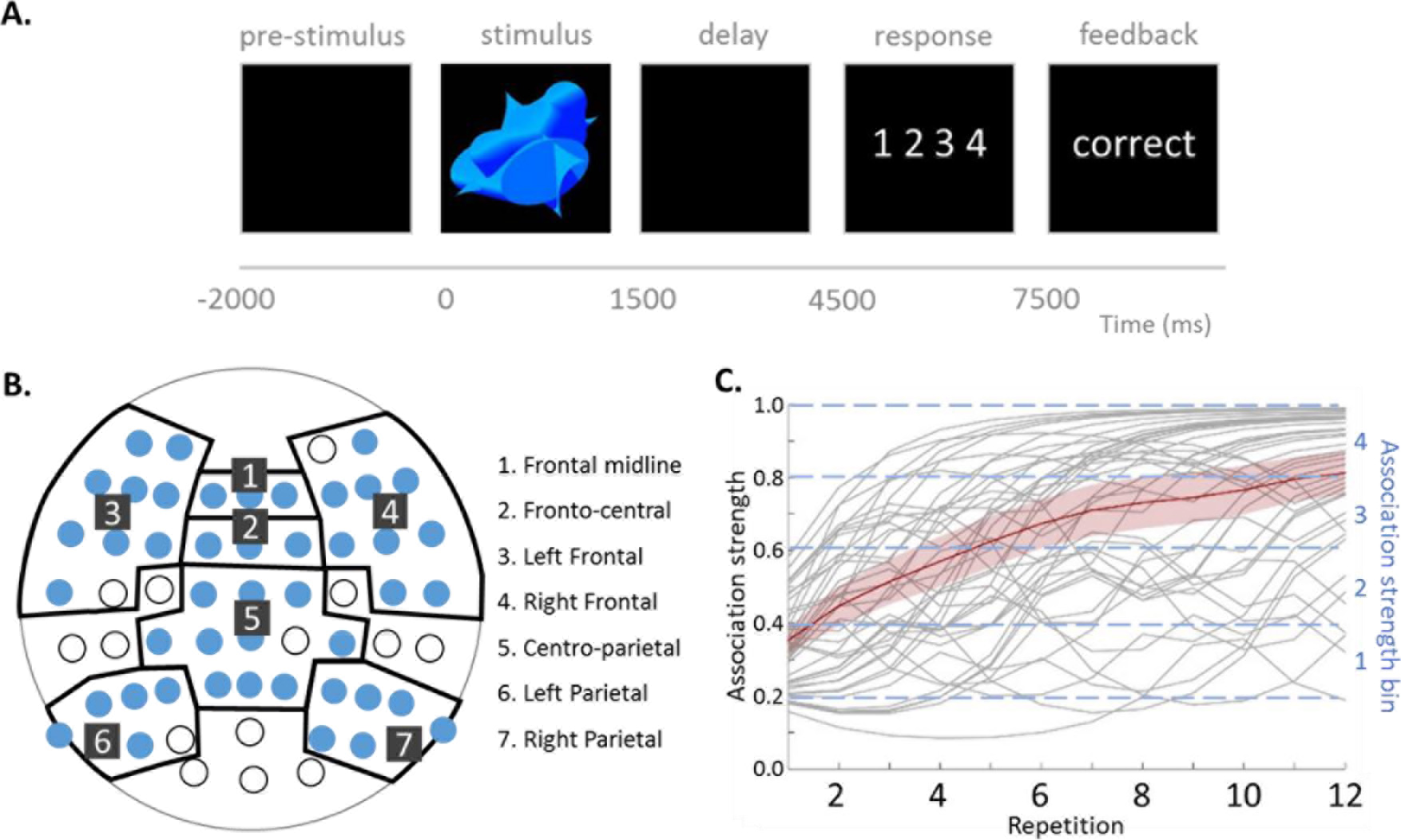
Experimental Approach. A. Example of a trial showing the task timings. B. Hierarchical clustering was used to group of electrodes into bilateral frontal, frontal midline, fronto-central, centro-parietal and bilateral parietal clusters. Unfilled circles show electrodes that did not cluster into this scheme or were their own cluster. C. Learning data for a representative subject. Individual learning curves for each stimulus-response association (n=48) are shown in grey, plotting the estimated association strength across repetitions. Red curve shows the average learning curve over trials and the standard error.

### EEG analysis

Preprocessing of the EEG used the EEGLAB toolbox (Delorme & Makeig, 2004) in Matlab. Data were referenced to the average of the left and right mastoids, highpass filtered at 0.5 Hz using an FIR filter of length 12288 points, and resampled to 500 Hz. Bad channels were identified by visual inspection and reconstructed using spherical interpolation. Data were epoched between −2 seconds and 8.5 seconds after the onset of the shape image, and baseline corrected using the −200 to 0 ms period. Independent component analysis (ICA) was performed using runica (Delorme, Sejnowski, & Makeig, 2007). SASICA and ADJUST (Chaumon, Bishop, & Busch, 2015; Mognon, Jovicich, Bruzzone, & Buiatti, 2011) were used for the detection of artifactual components to reject, which were validated through visual inspection (as recommended Chaumon et al., 2015). The data were transformed to a scalp surface Laplacian, or current source density estimate using the CSD toolbox (Kayser & Tenke, 2006). This is a reference-free estimate of the scalp current density, that minimises the effect of volume conduction to increase the spatial localisation relative to electrical scalp potentials (Nunez & Srinivasan, 2006). The current source density information was calculated using a smoothing constant of *lambda* = 1.0^−5^, head radius of 10 cm, and spline interpolation constant of *m* = 4. Time-frequency representations (TFRs) of oscillatory power between 4 and 60 Hz (in 30 log-spaced steps) were calculated for each trial using Morlet wavelets with a minimum 5-cycles increasing to a maximum of 15-cycles at 60 Hz. Oscillatory power was calculated between-1.25 seconds to 7.5 seconds in 50 ms steps (181 time points). Baseline correction was applied to each trial using a prestimulus period between −1.25 and −0.75 seconds.

### Electrode regions

Our analysis was performed across different electrode regions of interest. This allows us to reduce the number of statistical comparisons made, while allowing us to focus on bilateral frontal, frontal midline, fronto-central, centro-parietal and bilateral parietal electrode sites. Rather than arbitrarily grouping electrodes into clusters, we used a more data-driven approach to determine which electrodes should be grouped together with the proviso that we end up with bilateral frontal, frontal midline, fronto-central, centro-parietal and bilateral parietal regions. By using hierarchical clustering, we can determine which electrodes have similar oscillatory activity and determine how electrode regions are formed.

To determine electrode regions, TFRs were averaged across all trials and subjects to produce a grand-average for each electrode. Data for each electrode were vectorised, before hierarchical clustering of electrodes using correlation as the distance measure. The resulting distances were visualised as a dendrogram to define the initial state of the electrode regions. As shown in Figure 1B, hierarchical clustering showed 7 clusters that met our *a priori* scheme creating electrode regions in lateral frontal, parietal, central, and mid-frontal regions. Electrodes and clusters that did not fit into this scheme were excluded (e.g. AF4 was in a cluster separate to all other electrodes). Once the electrode regions were defined, single trial TFRs were averaged across electrodes within each region, to give seven TFRs for each trial that were tested against the learning data.

### Defining association strength

To test the relationship between learning and oscillatory power we need to define a quantitative measure of how much the association has been learned for any given trial. Following Smith et al., (2004), the binary performance data on each trial (correct/incorrect) were used to estimate a trial-wise measure of association strength, where for each trial we obtained a probability of correct response based on the previous responses to that trial type. Trial-specific learning curves were calculated (n = 48) for each subject, providing a quantitative measure of how strongly the association had been learned. Following Law et al., (2005), the continuous probability values from the learning curves were binned where association strength index 1 trials had a probability between 0.2-0.4, association strength 2 index trials had a probability between 0.4-0.6, association strength index 3 trials had a probability between 0.6-0.8, association strength index 4 trials had a probability between 0.8-1. Trials with association strength values less than 0.2 were discarded from all analyses as they reflect below chance performance (Figure 1C).

### Linear modelling

To test the relationship between EEG oscillations and association strength, linear fixed effects models were calculated for each subject, time, frequency point, and electrode region separately. Trials were excluded where the response was made prior to the response cue, and we excluded the first repetition of each item so that our effects could not be driven by a novelty effect. We also excluded individual time/frequency data-points from the linear modelling to ensure the analysis was not driven by outliers. At a given electrode, time, and frequency point, EEG data points that were more than 2 standard deviations away from the mean EEG response were excluded. The remaining EEG signals were the dependent variable, and predictor variables were trial number (a proxy for experimental time), the response time, previous trial response time, and the association strength index for that trial. This resulted in a beta-coefficient TFR for each subject, region and predictor variable that captured linear changes between association strength and EEG. Random effects analysis testing for positive or negative coefficients was conducted for each time-frequency point using one-sample t-tests against zero (alpha 0.05, two-tailed). Cluster-mass permutation testing was used to assign p-values to clusters of significant tests (Maris & Oostenveld, 2007), and a maximum cluster approach was used to control for multiple comparisons across time, frequency and electrode regions (Nichols & Holmes, 2002). For each permutation, the sign of the beta-coefficient was randomly flipped for each subject before one-sample t-tests of the permuted data. The same permutation was applied to all electrode regions, and the cluster with the largest mass across all regions (sum of t-values) was retained. The p-value for each cluster in the original data was defined as the proportion of the 10,000 permutation cluster-masses (plus the observed cluster-mass) that is greater than or equal to the observed cluster-mass.

## Results-Behaviour

Participants were highly successful at learning each association. On average, 90% of stimulus-response associations were correct on the final repetition (min 71%, max 100%). In the final testing session, conducted after the final block, an average of 84% of stimulus response associations were correctly answered (min 67%, max 100%).

Conditional associative learning is often characterized as a gradual process, but as noted by Gallistel (2004), this impression can be an artefact of averaging learning rates across individual associations. In fact, as shown in Figure 1C, learning curves for individual associations within a subject are highly variable, often showing abrupt transitions resembling a sigmoidal function. Accordingly, to characterize learning in this task, it is essential to quantify memory for individual associations rather than average learning curves. To obtain learning curves for individual associations, we used the state-space model introduced by Smith et al. (2004). This modelling approach allowed us to estimate the strength of learned associations on a trial-wise basis. This continuous measure was then transformed into four association strength bins (0.2-0.4, 0.4-0.6, 0.6-0.8, 0.8-1.0). Trials with association strengths between 0 and 0.2 were excluded from all analyses as they were well below chance (i.e., less than 25% accuracy) performance. Using this approach, for each presentation of a stimulus, we obtained measures of association strength that could be tested against single-trial data.

A linear mixed effects model tested the relationship between association strength and reaction time. Results showed that association strength was inversely related to reaction times (β = −38 ms, t(6775) = −2.26, p = 0.024). We further included nuisance variables in the model to account for effects of fatigue, attention/arousal and experience with the task. Reaction time was significantly related to trial number (β = −0.11 ms, t(6775) = −5.96, p <0.0001), and reaction time on the previous trial (β = 0.01 ms, t(6775) = 2.84, p = 0.005). These results show that stronger visuo-motor associations have faster reaction times, an effect that is over and above the impact of nuisance factors such as trial number. The relationship between reaction time and association strength highlights the importance of taking into account reaction time, and other nuisance measures, in our EEG analysis, to ensure the results are not confounded.

## Results-EEG

### Association strength

Our primary goal was to test how oscillations change in relation to memory performance. Specifically, we tested for a linear relationship between association strength and oscillatory power, while controlling for nuisance effects of experimental time and overall reaction time. As shown in Figure 2, this analysis revealed significant effects in theta at frontal midline sites during the delay period, and posterior alpha and beta effects during the cue, delay and response periods. We further see a right frontal effect in alpha and beta around the onset of the response period.

**Figurre 2.**
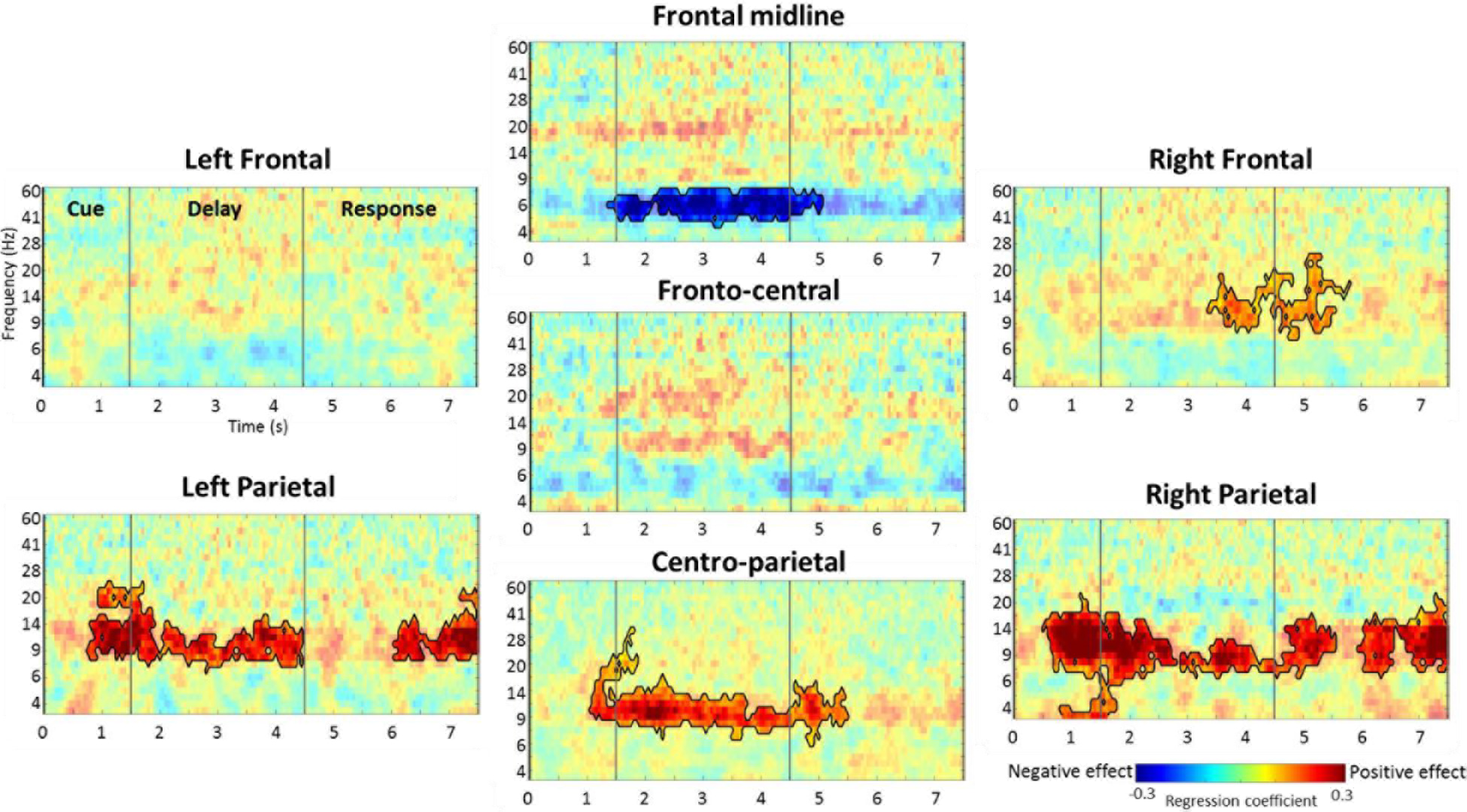
Overview of the results. Frontal midline theta shows a negative relationship with increasing association strength, while posterior alpha/beta, and right frontal, show positive relationships with increasing association strength. Time-frequency spectograms show the regression coefficients from a linear model between oscillatory power and association strength, while controlling for nuisance variables. Vertical lines show the divisions between the cue, delay and response periods. Significant effects are outlined in black.

We first concentrate on effects of association strength on theta power (Figure 3). Frontal midline electrode sites showed a significant negative linear effect of association strength that largely overlapped with the delay period (5-8 Hz, 1350 to 5050 ms; cluster p = 0.004). By examining the overall mean oscillatory activity across trials (Figure 3A, left), we can see that there is overall theta synchronisation during the delay period (compared to a pre-stimulus baseline period), and our finding of a negative effect of association strength shows that the amount of theta synchronisation reduces as the associations are learned. Plotting the mean theta activity for each association strength bin illustrates this linear reduction in theta activity as associations are learned (Figure 3A, right), which is also evidenced by plotting the oscillatory activity for each association strength bin (Figure 3B). These changes in theta activity suggest that frontal midline sites support associative learning, with theta rhythms important during the delay period when stimulus information and decisions are maintained, and that as associations are learned theta activity decreases.

**Figure 3.**
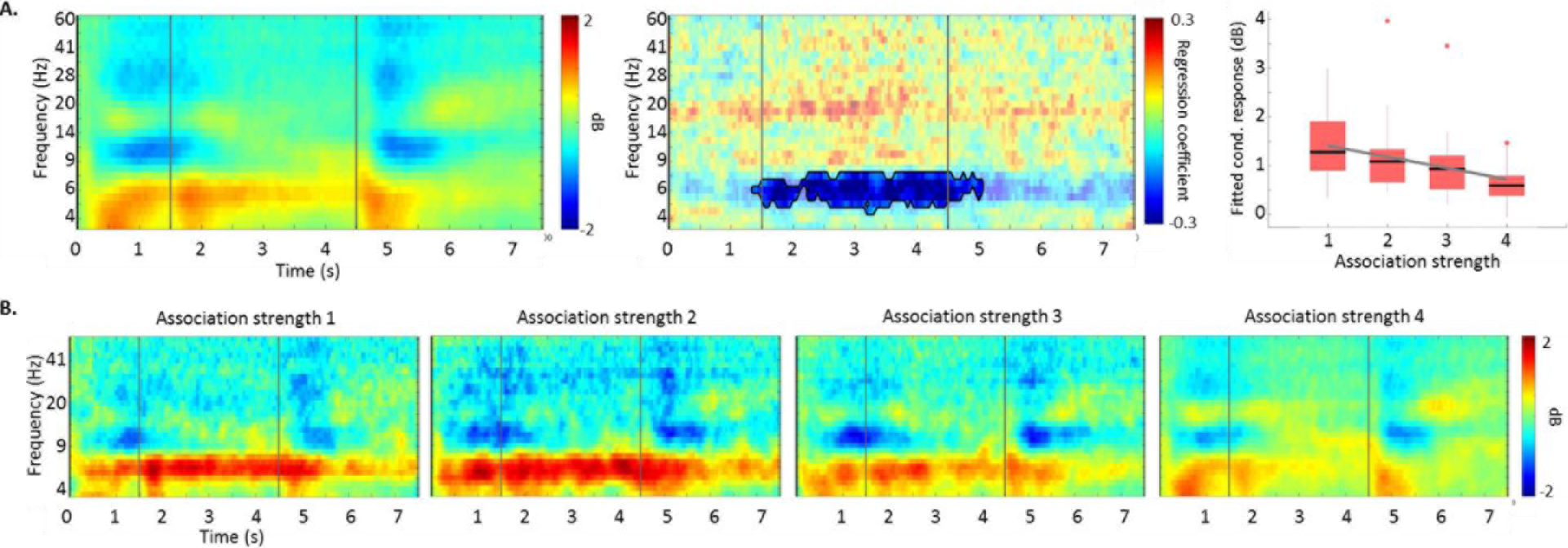
Frontal midline theta decreases with increasing association strength. A) Left, time-frequency spectogram showing the mean power over all trials compared to a pre-stimulus baseline period. Vertical lines show the divisions between the cue, delay and response periods. Middle, regression coefficients from a linear model between power and association strength showing negative effects during the delay period. Significant effects are outlined in black. Right, plot showing the relationship between association strength bin and power from the significant cluster. Trend line shows the linear relationship between association strength and power, while boxplots show the distribution of individual participant data (outliers shown as dots). B) Time-frequency spectograms for each association strength bin to show how power is greatest for association strength bin 1 and 2 and decreases towards baseline levels as the associations are learned.

We next focused on the significant relationships between associative strength and alpha and beta power. We found a positive relationship between association strength and oscillatory activity spread across alpha and beta frequencies in left, right and centro parietal electrode sites (Figure 4), and in the right frontal region. In the left parietal region, significant linear effects were found in two clusters, between 7-24 Hz and 750 to 4450 ms (p = 0.002), and between 8-22 Hz and 6050 to 7500 ms (p = 0.026). The centro-parietal region showed a significant linear effect between 6-31 Hz and 1050 to 5500 ms (p = 0.001) although the focus of this effect is within alpha frequencies during the delay period. Three significant linear clusters were found in the right parietal region; one focussed on the cue-delay period between 4-22 Hz and 500 to 2800 ms (p = 0.001), one on the delay-response period between 7-16 Hz and 2900 to 5550 ms (p = 0.016), and one during the response period between 7-22 Hz and 5950 to 7500 ms (p = 0.014).

**Figure 4.**
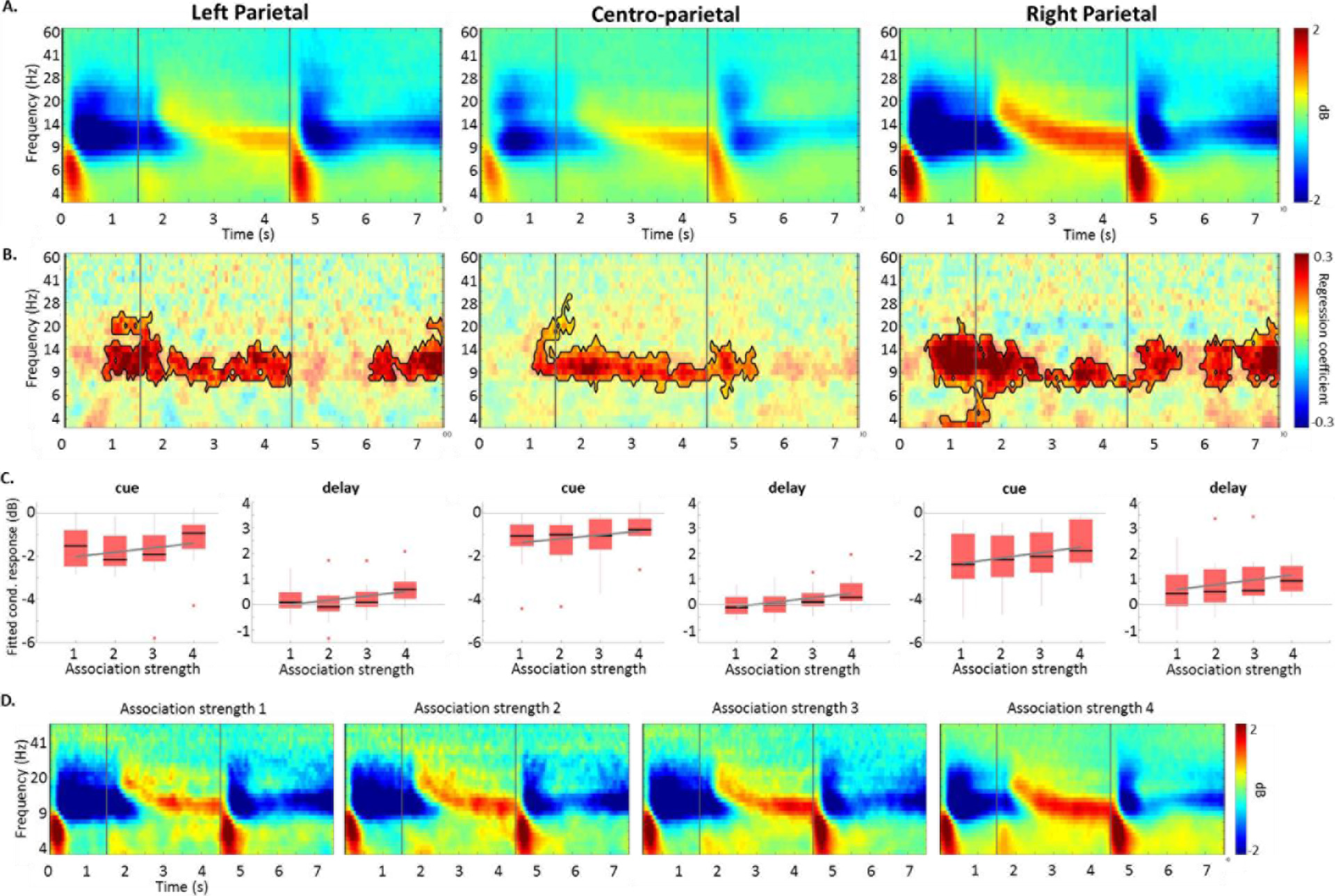
Increases in alpha and beta power with increasing association strength. A) Time-frequency spectograms showing the mean power over all trials compared to a pre-stimulus baseline period. Vertical lines show the divisions between the cue, delay and response periods. B) Regression coefficients from the linear model between oscillatory power and association strength showing positive effects. Significant effects are outlined in black. C) Plots showing the relationship between association strength bin and power from the individual significant clusters, separated into the cue and delay period. Trend lines show the linear relationship between association strength and power, while boxplots show the distribution of individual participant data. D) Time-frequency spectograms from the right parietal sites for each association strength bin to show how cue period alpha/beta desynchronisation is greatest for association strength bin 1 and 2 and decreases as the associations are learned, which delay period alpha synchrony increases with association strength.

When considering overall oscillatory activity over all trials, bilateral parietal and centro-parietal electrode sites exhibit alpha and low beta desynchronisations in the cue and response period, and alpha synchronisations during the delay period (Figure 4A,D). Plotting the oscillatory activity from each significant cluster, in the cue and delay periods separately, shows that as association strength increases, there is less cue period desynchronization (less negative power compared to a prestimulus baseline; Figure 4C) while there is more delay period alpha synchronisation (i.e. overall alpha power is positive and gets more positive as the associations are learned). This shows that cue period desynchronisations are greatest during early phases of learning and reduce as the associations become well learned, while delay period alpha synchronisations increase as the associations are learned (Figure 4D). Right frontal electrode sites also showed a significant positive linear effect of association strength (8-24 Hz, 3350 to 5800 ms; p = 0.020) that is spread across the delay-response period.

### Learning effects

The above analyses demonstrate that theta power is high early in the associative learning process and decreases during learning. In addition, cue period alpha and beta oscillations show large desynchronisations early in learning that reduce as the associations are formed, and delay period alpha synchronisations increase with learning. This relationship between association strength and oscillatory activity could reflect learning processes, or processes that are likely to be differentially engaged during the learning process (e.g. cognitive control processes such as working memory demands and response uncertainty). Accordingly, we ran a second analysis to test whether oscillations during trials were predictive of how much learning will take place on that trial. Specifically, we tested for a “learning effect,” operationalized as a linear relationship between oscillatory power on the current trial and the change in association strength from the current trial to the next presentation. This analysis can be seen as analogous to the kinds of “difference memory” or “subsequent memory effect” analyses that are used in studies of single-trial learning.

Learning effects were defined as the difference in association strength between a trial and its next presentation. We chose to limit our analysis to trials that showed chance performance-those in association strength bin 1, as they provide the maximal opportunity for, and variability in, learning, and all trials should require similar cognitive control processes such as working memory demands and response uncertainty. Rather than calculate the difference in association strength using the association strength bins, the difference in association strength was based on the original continuous measure from the state-space model of learning. Once learning effects were calculated, these values were divided into 5 bins capturing when learning effects were negative (i.e. when the difference in association strength was less than 0; bins-2 and-1), and when learning effects were positive (i.e. when the difference in association strength was greater than 0; bins 1, 2 and 3). The range of association strength values in each bin was 0.2.

Learning effects were tested for data averaged across the significant time-frequency points reported for the association strength analysis above. As in the analyses described above, we used linear fixed effects models and controlled for nuisance effects of experimental time and overall reaction time. Frontal midline theta showed a significant positive relationship where power was greater for items showing larger learning effects (β = 0.20, t(15) = 2.22, p = 0.042; Figure 5). Significant negative relationships between power and learning effects were observed for right parietal and right frontal, where greater learning was associated with increasing desynchronisations in alpha and low beta frequencies (Right parietal: cue/delay β = −0.15, t(15) = −2.22, p = 0.042; delay/response β = −0.24, t(15) = −2.53, p = 0.023; late response β = −0.27, t(15) = −4.77, p = 0.0002. Right frontal: delay/response β= −0.14, t(15)= −2.61, p =0.020). Follow-up analysis of the right parietal cluster, showed significant effects were present when the analysis was restricted to the cue period (β = −0.17, t(15)=-2.32, p = 0.035) but only trended in the delay period (β = −0.21, t(15)=-1.9, p = 0.072). These results show that the oscillatory responses that were modulated by association strength, were further predictive of the amount of learning that will occur, showing that these low-frequency oscillations are important for memory formation.

**Figure 5.**
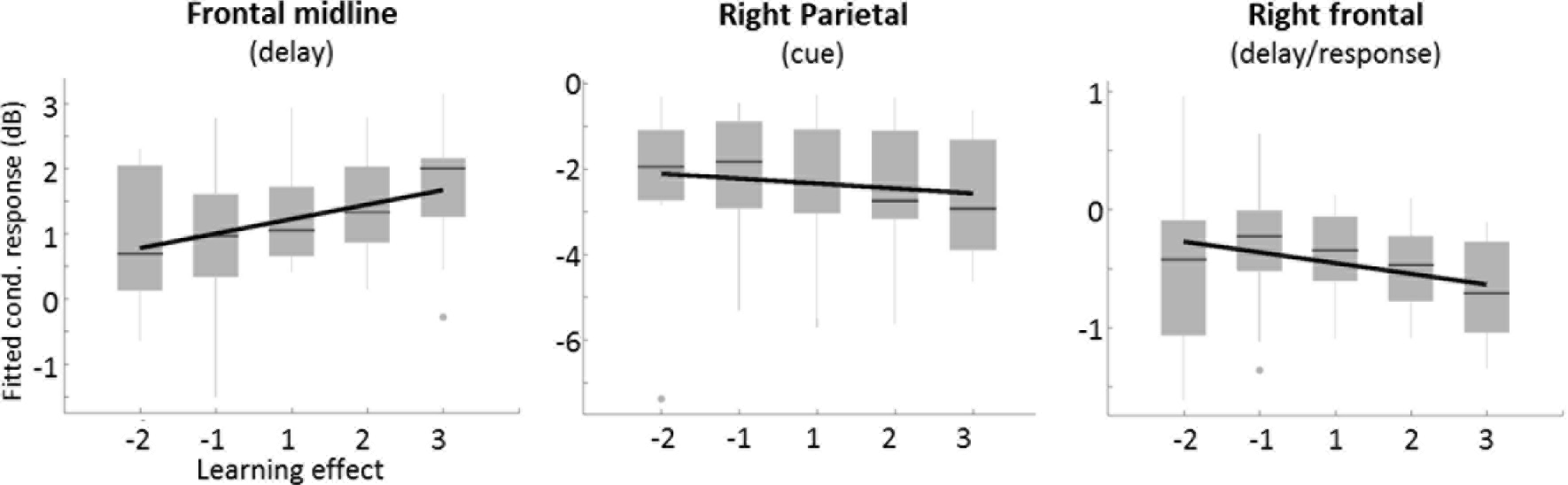
Relationship between oscillatory power and learning effects. Learning effects are shown on the x-axis where negative values show a decrease in association strength on the subsequent trial, and positive values show increasing changes in association strength for the subsequent trial. Trend lines show the linear relationship between learning effects and oscillatory power, while boxplots show the distribution of individual participant data.

## Discussion

The goal of the present study was to examine how neural oscillations change during the learning of arbitrary visuo-motor associations in terms of (1) how oscillations change as memories get stronger, and (2) whether oscillations are directly related to learning. By differentially relating single-trial measures of association strength to neural oscillations – while controlling for other factors that coincide with learning such as time, we showed that theta, alpha and beta oscillations track associative memory formation, and that these effects predicted increments in associative memory strength on the next occurrence. We show that cue period alpha and beta desynchronisations are greatest early in learning, coupled with delay period frontal midline theta. Both alpha/beta desynchronisations and theta synchronisations reduced as the associations became well learned. These early alpha/beta desynchronisations and theta synchronisations further correlated with how much learning would occur on a given trial, as shown by our analysis of learning effects. In addition, we show that as associations become well learned, delay period alpha synchronisations increased.

Our finding that neural oscillations show a linear relationship with association strength mirrors other research that used state-space models to characterise learning and memory formation (Hargreaves et al., 2012; Law et al., 2005; Wirth et al., 2003). For example, using fMRI, Law et al., (2005) used a similar associative learning paradigm to the one employed here, and found that BOLD signals in the MTL, frontal and parietal regions showed a linear relationship with association strength. Like in the present study, Law et al., (2005) showed both positive and negative relationships with association strength. They report that the MTL and medial frontal regions show a positive relationship, while parietal and frontal regions showed a negative relationship. In the present study, frontal theta power showed a negative relationship with association strength, and posterior sites showed a positive relationship (in addition to right frontal sites), showing an inverse relationship between the direction of frontal and parietal effects seen across the two studies. However, this can be reconciled by considering the inverse relationship between theta and BOLD that has been noted in medial PFC and ACC (Scheeringa et al., 2008)-regions thought to generate frontal midline theta oscillations (Hsieh & Ranganath, 2014; Onton, Delorme, & Makeig, 2005; Raghavachari et al., 2001; Tsujimoto, Shimazu, & Isomura, 2006), and show connectivity with the HPC during memory tasks (Brincat & Miller, 2015; Fuentemilla, Barnes, Düzel, & Levine, 2014; Garrido, Barnes, Kumaran, Maguire, & Dolan, 2015). This suggests that both our results, and those found in fMRI, capture the same underlying process during memory formation, with the additional temporal and spectral resolution here across the cue and delay periods.

Our results show that delay period frontal midline theta plays a prominent role during associative memory formation. This was shown across two analyses that tested the relationship between oscillations and estimates of associative memory strength. We found maximal frontal midline theta power was observed during poorer association strength trials, during the early learning of the association, which decreased as association strength increased. Further, during trials that had chance memory performance, the degree of theta activity related to memory improvements when the item was next seen (an analysis similar in nature to a subsequent memory, or difference memory effect), where more theta power signalled a greater improvement in memory. Together, these results show that frontal theta activity enhances the learning of visuo-motor associations.

Theta oscillations in humans have been linked to a wide range of memory phenomena including memory encoding (Greenberg, Burke, Haque, Kahana, & Zaghloul, 2015; W. Klimesch et al., 1997; Mölle et al., 2002; Olsen, Rondina, Riggs, Meltzer, & Ryan, 2013; Rutishauser, Ross, Mamelak, & Schuman, 2010; Sederberg, Kahana, Howard, Donner, & Madsen, 2003; Staudigl & Hanslmayr, 2013; Summerfield & Mangels, 2005), retrieval (Addante, Watrous, Yonelinas, Ekstrom, & Ranganath, 2011; Burgess & Gruzelier, 1997; Gruber, Tsivilis, Giabbiconi, & Müller, 2008; Guderian & Düzel, 2005; Jacobs, Hwang, Curran, & Kahana, 2006; W. Klimesch et al., 2001) and working memory maintenance (Gevins, Smith, McEvoy, & Yu, 1997; Hsieh, Ekstrom, & Ranganath, 2011; Jensen & Tesche, 2002; Olsen et al., 2013; Roberts, Hsieh, & Ranganath, 2013), underlining its central role in long term memory. Our theta effects of association strength were found during the delay period, although overall theta power increased during the cue period and remained elevated throughout the delay until a response was made (Figure 3B). This sustained theta activity could reflect the maintenance of information in working memory, and echoes recordings from the human middle frontal gyrus where theta power increases during working memory maintenance (Raghavachari et al., 2001). This suggests that higher theta activity early in learning could reflect better maintenance of cue information, that in turn supports better learning of the cue-response association.

This is supported by previous EEG research that explored the relationship between neural oscillations during active maintenance of objects and successful long-term memory encoding. Khader et al., (2010) found subsequent memory effects of theta oscillations during a delay period, in that theta activity was greater for successfully encoded items compared to items that were later forgotten. Here, we show delay period theta activity was greatest during early learning and reduced as the association was learned. In addition, our analysis of learning effects showed that theta activity early in learning was greater for trials that showed the greatest improvements in association strength, an analysis similar in nature to a subsequent memory effect. Therefore, our theta effects during the delay period likely reflect the maintenance of cue information, that in turn leads to better encoding of the cue item and learning of the cue-response association. This would be especially important during early learning. Late in learning, delay period theta activity decreases to near baseline levels as the associations are well learned.

We further saw a modulation of alpha and beta activity as associations were formed in both the cue and delay periods over posterior electrodes. During the cue period, alpha and beta desynchronisations (defined as lower power compared to a pre-stimulus period) were greatest for low association strength trials and tended back towards a baseline level as associations became well learned. Posterior alpha and beta desynchronisations are thought to reflect the active engagement of regions, in contrast to posterior alpha synchronisations thought to reflect the inhibition of task-irrelevant regions (Jensen & Mazaheri, 2010). Alpha and beta desynchronisations are claimed to play an active role in memory formation (Hanslmayr et al., 2012), although this has largely been reported in relation to the encoding of single items. Greater alpha desynchronization during stimulus presentation has been reported to predict better encoding of items (Hanslmayr et al., 2009; W. Klimesch et al., 1996, 1997; Mölle et al., 2002), and here we show the role of alpha and beta power decreases in memory formation extends to associative memory. Cue period alpha and beta desynchronisations were greatest during early learning, and further related to memory improvements when the item was next seen. This active engagement of posterior sites during the cue period likely reflects the active encoding and processing of stimulus information that supports memory formation, which would further support the maintenance of stimulus information through theta activity in the delay period.

The combination of theta synchronisation and alpha desynchronization may be a salient feature of successful memory formation (Hanslmayr et al., 2012; Parish, Hanslmayr, & Bowman, 2017). Frontal theta has been reported to show an inverse relationship to posterior alpha during the maintenance of temporal order information in working memory (Hsieh et al., 2011), while other research has shown that conjoint theta synchronisations and alpha desynchronisations support the successful encoding of items (Mölle et al., 2002). Together, we suggest that early in learning, alpha and theta oscillations together play a crucial role in the processing and maintenance of stimulus information that leads to better encoding of the stimulus and learning of the cue-response association.

We also observed a positive relationship between alpha synchronisation and association strength during the delay period, in that alpha activity increased with learning. According to the gating-by-inhibition hypothesis, alpha synchronisation over posterior sites reflects the inhibition of task-irrelevant regions (Jensen & Mazaheri, 2010; Wolfgang Klimesch, Sauseng, & Hanslmayr, 2007), suggesting that as the associations become well learned, posterior sites are increasingly inhibited during the delay period. Previous research has found that greater alpha activity during a maintenance period predicts the successful encoding of item information (Khader et al., 2010; Meeuwissen, Takashima, Fernández, & Jensen, 2011). This is interpreted within a framework where alpha activity during the maintenance of information in working memory reflects the inhibition of sensory, or bottom up areas, leading to better internal cognitive processing and memory encoding (Jensen, Gelfand, Kounios, & Lisman, 2002; Jensen & Mazaheri, 2010; Wolfgang Klimesch et al., 2007). Our results show that delay period alpha activity increases as the associations are learned, and so could reflect the increasing inhibition of task-irrelevant posterior sites to enhance memory retrieval of the correct cue-response associations.

This also highlights two dissociable functions of alpha oscillations at different timeframes. Early in learning, alpha desynchronisations during the cue period are important for the active encoding of the stimulus and successful learning. Late in learning, alpha synchronisations during the delay period could aid successful retrieval by inhibiting posterior sites related to sensory processing. Previously, both alpha synchronisations and desynchroniations have been shown to relate to memory performance across different studies (Hanslmayr et al., 2009; Khader et al., 2010; W. Klimesch et al., 1996, 1997; Meeuwissen et al., 2011; Mölle et al., 2002). These differences have been attributed to either an active encoding of stimulus information through desynchronisations or the inhibition of task-irrelevant areas during maintenance (Hanslmayr et al., 2012), with our study highlighting the duel effects of alpha within a single study.

## Conclusions

Learning is a dynamic process where neural representations are refined and change with experience. Our results highlight a central role of frontal theta and posterior alpha/beta oscillations in forming new associative memories. By relating neural oscillations to single-trial measures of associative memory strength - derived from state-space computational models of learning, we found theta and alpha/beta oscillations tracked learning behaviour. Early in learning, we find cue period alpha desynchronisation and delay period theta synchronisation are greatest and decline as associations are formed. These alpha and theta effects reflect the engagement of posterior regions for stimulus processing and encoding, and theta supports better maintenance of stimulus information. Further, the greater the alpha and theta effects are early in learning, the more of an increase in association strength we see. As the associations become well learned, we see increased alpha synchronisations during the delay period showing the inhibition of posterior regions. Our results suggest differential roles of alpha and theta during early learning, relating to stimulus encoding and maintenance, and different roles for alpha oscillations over the course of learning – moving from encoding of stimulus properties to inhibition of sensory regions when they are no longer task-relevant. These results provide direction for future studies, that can manipulate frontal theta and posterior alpha/beta oscillations to enhance learning efficacy and speed using non-invasive brain stimulation techniques.

## Acknowledgements

This work was supported by a Vanneavar Bush Faculty Fellowship (Office of Naval Research Grant N00014- 15- 1-0033) to CR. Any opinions, findings, and conclusions or recommendations expressed in this material are those of the author(s) and do not necessarily reflect the views of the Office of Naval Research or the U.S. Department of Defense.

